# Death and population dynamics affect mutation rate estimates and evolvability under stress in bacteria

**DOI:** 10.1101/224675

**Authors:** Antoine Frénoy, Sebastian Bonhoeffer

**Affiliations:** Institute for Integrative Biology, ETH Zürich, Switzerland

## Abstract

The stress-induced mutagenesis paradigm postulates that in response to stress, bacteria increase their genome-wide mutation rate, in turn increasing the chances that a descendant is able to withstand the stress. This has implications for antibiotic treatment: exposure to sub-inhibitory doses of antibiotics has been reported to increase bacterial mutation rates, and thus probably the rate at which resistance mutations appear and lead to treatment failure.

Measuring mutation rates under stress, however, is problematic, because existing methods assume there is no death. Yet sub-inhibitory stress levels may induce a substantial death rate. Death events need to be compensated by extra replication to reach a given population size, thus giving more opportunities to acquire mutations. We show that ignoring death leads to a systematic overestimation of mutation rates under stress.

We developed a system using plasmid segregation to measure death and growth rates simultaneously in bacterial populations. We use it to replicate classical experiments reporting antibiotic-induced mutagenesis. We found that a substantial death rate occurs at the tested sub-inhibitory concentrations, and taking this death into account lowers and sometimes removes the signal for stress-induced mutagenesis. Moreover even when antibiotics increase mutation rate, sub-inhibitory treatments do not increase genetic diversity and evolvability, again because of effects of the antibiotics on population dynamics.

Beside showing that population dynamic is a crucial but neglected parameter affecting evolvability, we provide better experimental and computational tools to study evolvability under stress, leading to a re-assessment of the magnitude and significance of the stress-induced mutagenesis paradigm.

## Introduction

One of the most puzzling and controversial microbial evolution experiments of the 20th century may be the one performed by Cairns and collaborators [8, 7] in which *lac*– cells are plated on lactose as the sole carbon source and therefore cannot grow. Revertants toward the lac+ genotype continuously appear after plating at a rate and timing seemingly incompatible with the Darwinian hypothesis of selection of pre-existing mutants. In the *lac*– construct, the *lacZ* coding sequence is present but non-functional, because it is out of frame with the start codon. The *lac*+ revertants are thus frameshift mutants in which this coding sequence is back in frame with the start codon. Most of the controversy initially came from the question of whether these reversion mutations where Lamarckian, in the sense that they would arise at a higher rate when the cells would “sense” that these mutations would be beneficial [45]. However many additional experiments quickly suggested that this phenomenon can be explained by more standard Darwinian mechanisms, where genetic changes are not targeted, but occur randomly and are then selected or not. While two seemingly conflicting molecular explanations emerged — the stress-induced mutagenesis model and the gene amplification model —, both are conceptually very similar.

In both explanations, mutations occur randomly and independently of their effect on fitness, but the specific conditions of carbon starvation increase the rate at which genetic diversity is generated at the relevant locus (*lacI-lacZ* sequence). In the stress-induced mutagenesis model [53], the genome-wide mutation rate is increased as an effect of the stress response triggered by starvation. In the gene amplification model [2], random duplications of the *lacI-lacZ* system happen and are selected because the frameshift mutation is leaky. A small amount of Beta-galactosidase is still synthesised permitting cryptic growth due to rare expression errors, which compensate the frameshift. This residual expression becomes bigger with more copies of the leaky system. From an initial duplication a further increase in copy number will be favored by recombination due to sequence homology. As the copy number of the system increases, a reversion mutation in *lacI-lacZ* becomes more likely, because of increased target number.

While it is still not clear whether stress-induced mutagenesis is the sole explanation of the phenomenon, the attempts to explain the data presented by Cairns and collaborators have lead to a much better understanding of control over mutation rate in response to the environment. Such increase of mutation rate under starvation has also been reported in other systems [46, 4, 35]. An emblematic molecular mechanism permitting this regulation of mutation rate is the SOS response, suggested in 1970 [38, 42], in which DNA damages are sensed by bacterial cells and lead to the up-regulation of many genes permitting mutagenic repair and replication of damaged DNA. While the responsible enzymes were unknown at the time, it has indeed been found subsequently that the SOS response increases the dosage of polIV and polV [12]. These error prone polymerases are able to replicate damaged DNA that the classical DNA polymerase polIII could not replicate, albeit at the price of a higher error rate [56, 49]. This strategy favoring “survival at the price of the mutation” is only one side of the story. There is a line of evidence suggesting that this higher error rate is not only an unavoidable trade-off with survival. It is also supposed to be a selected property to increase mutation rate under stressful conditions, increasing the chances that one of the descendants obtains a beneficial mutation, which makes it able to better withstand the stress [20, 39].

The evolution of traits that increase mutation rate under stress needs be considered in the context of second-order selection [51]. Second-order selection relies on the idea that natural selection does not only act on the individual’s phenotype and instant fitness, but also on its ability to generate fit descendants, leading to selection of properties such as evolvability and mutational robustness [17]. In parallel to the study of environmental control over the mutation rate, genetic determinants of mutation rate have also been studied. It has been shown and is widely accepted that alleles increasing mutation rate, for example defective mismatch-repair or DNA proofreading, can be selected for the resulting increase in evolvability [9, 47]. On the other hand the possibility of selection of mechanisms increasing mutation rate under stress but not constitutively has been subject to a more philosophical debate [10, 43]. While modeling shows such selection is possible [39], it is hard to distinguish whether an observed increase in mutation rate under a specific stress is (i) an evolvability strategy, (ii) an unavoidable trade-off of selection for survival, such as replicating damaged DNA to avoid death at the price of making mutations, or (iii) a direct effect of the stress and not of the stress-response system [34].

But this debate does not affect the evolutionary relevance of the phenomenon, nor the medical implications concerning the risk of de novo evolution of resistance during antimicrobial treatment. Here we are interested in the general case of mutation rate in growing stressed populations, and we especially focus on antibiotic stress, although our findings may be valid for many other biotic and abiotic stresses. It has been suggested that treatment with sub-inhibitory doses of antibiotics increases bacterial mutation rate, due to induction of various stress-response pathways [28, 3, 25, 41, 32]. Many molecular mechanisms underlying this stress response have been elucidated, including the SOS response [3] or the RpoS regulon [25]. Oxidative damages have also been suggested to play a role in antibiotic-induced mutagenesis [28] and death [16]. Although still controversial [31] these findings link antibiotic stress to the older question of how bacteria deal with oxidative stress and how oxidative damages impact mutation rates [36].

However all the evidence for stress-induced mutagenesis relies on accurately measuring mutation rates of bacteria growing in stressful conditions, and comparing them to those of the same strains growing without stress. Computing such mutation rates under stress is harder than it may seem, because stress may change population dynamics and may thus invalidate the hypotheses made by the mathematical models used to compute mutation rate. For example in the case of sub-inhibitory concentrations of antibiotics when net population growth is positive, death may nevertheless happen at a considerable rate. Death events, however, are not detected by standard microbiology methods, and are not taken into account by the mathematical tools used to compute mutation rate [59, 26, 22].

Such tools indeed only take as inputs the number of observed mutants at a chosen locus and the final population size, making the underlying assumption that there is no death and that population size is thus a sufficient information to summarise growth dynamics. The final population size is used to infer the number of DNA divisions leading to the final observed population from a small initial inoculum. If there is death, more divisions are needed to reach this population size, thus giving more opportunities to acquire mutations. The mutation rate will then be over-estimated, because the number of DNA replications will be under-estimated.

In this work we developed an experimental system to compute death rates in populations growing under stress, and a computational method to compute mutation rates from fluctuation assays under stress using the computed death rates. We applied this framework to re-estimate mutation rates of *Escherichia coli* MG1655 growing under sub-MIC doses of kanamycin (an aminoglycoside), norfloxacin (a fluoroquinolone) and hydrogen peroxide (an oxidizing agent). All these antimicrobials have previously been reported to significantly elevate mutation rate [28, 41]. For norfloxacin and kanamycin, we find that neglecting death leads to substantial overestimation of mutation rate. After conservatively correcting for death, the estimated increase of mutation rate due to treatment is largely reduced, and there remains no signal of stress-induced mutagenesis in the case of kanamycin.

We also show that mutation rate estimation does not only present experimental and mathematical challenges, but is also not the most relevant measure of evolvability meaning the capacity of a population to generate adaptive genetic diversity. Indeed, some of the studied sub-inhibitory treatments cause a significant drop in population size due to both bactericidal and bacteriostatic effects, and thus lead to a smaller genetic diversity despite a higher mutation rate. Ironically, evolvability can be much more easily estimated from experimental data than mutation rate. In our experiments, antibiotics and hydrogen peroxide have very different effects on evolvability: both sub-inhibitory norfloxacin and kanamycin treatments significantly reduce it, while hydrogen peroxide treatment strongly increases it.

## Results

### Mutation rates are over-estimated when neglecting death

Sub-inhibitory treatments are not necessarily sub-lethal, because minimal inhibitory concentration (MIC) is defined at population scale. An antimicrobial treatment is sub-inhibitory if the population grows (i.e. CFU/mL increases, or more crudely culture tubes inoculated at low density are turbid after 24h). However the death rate can be high, as long as the division rate is higher. Such death events will not be visible to the observer if only population size (CFU/mL) is tracked over time (figure 1). To reach a given, observed final population size, the number of cell divisions has to be higher, if there is death. This means that when computing mutation rate using the classical approach (described in the materials and methods), the number of cell divisions will be under-estimated. This is because it is implicitly assumed that there is no death and thus that the final population size is a good approximation for the number of cell divisions. The mutation rate, computed as the number of mutational events divided by the number of cell divisions, will then be systematically over-estimated.

**Figure 1:**
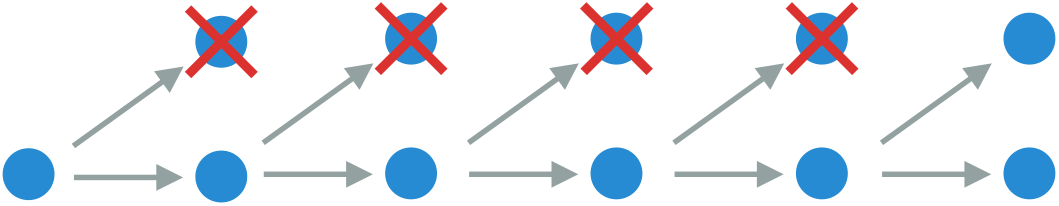
Example with death rate 0.8. One cell division ‘detected’ by change in population size requires actually five ‘real’ cell divisions. Each of these ‘hidden’ DNA replications gives extra chances to acquire a mutation.

The above statement, that mutation rates are systematically over-estimated when there is death, is the first intuition motivating our work. We explore this intuition more rigorously further below using a simulation approach. For an arbitrary chosen value of mutation rate toward a neutral arbitrary phenotype, we simulate the growth of a population of bacteria inoculated from a small number of non-mutant cells, with a chosen constant death rate, and track the number of mutant and non-mutant cells. We then compute the mutation rate based on the final state of these simulations, using the standard approach (i.e. the fluctuation test as described in the materials and methods) to test whether we recover the true value of the mutation rate. As shown in figure 2, the mutation rate is systematically over-estimated when there is death, and the higher the death rate the higher the over-estimation. This result is robust to changes in other population growth parameters such as the initial and the final population size, the mutation rate, and the plating fraction (data not shown).

**Figure 2:**
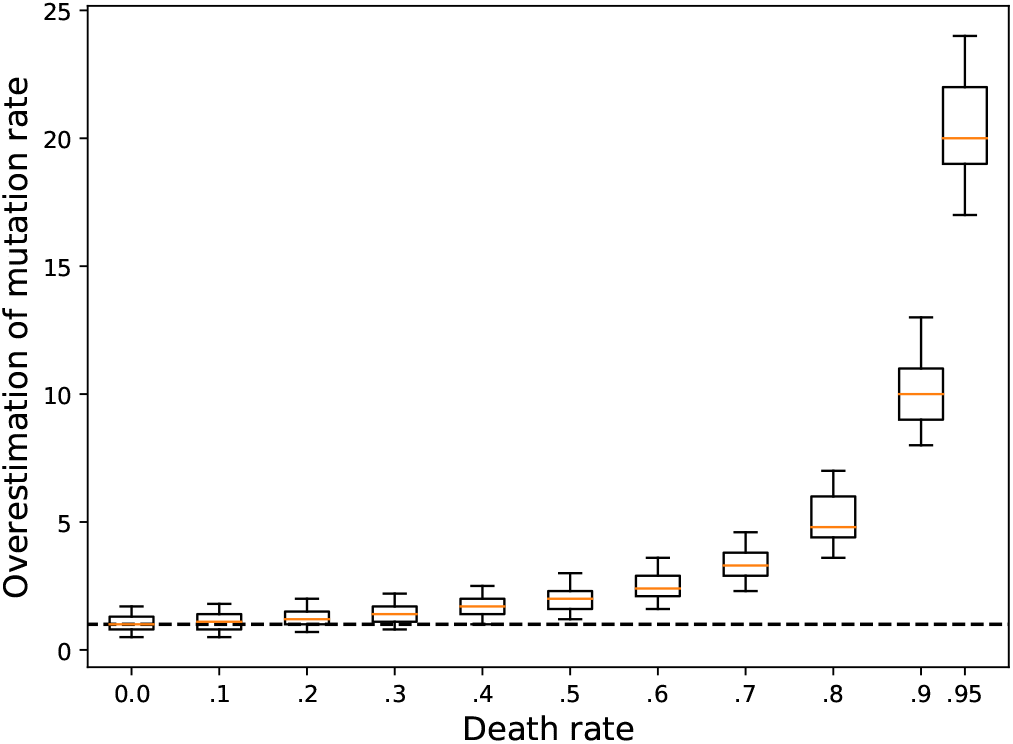
Over-estimation of mutation rate when there is death. From simulations of population growth with known death and mutation rate, we estimate the mutation rate using the classical method which does not take death into account. For each death rate between 0 and 0.95, 1000 simulations with 24 parallel cultures were performed. For each simulation we plot the ratio between the computed mutation rate (based on the number of mutants in the final state of the simulations) and the true mutation rate (used as input of the simulations, here 1 * 10^−9^). The orange lines indicate the median values, the boxes indicate the upper and lower quartiles, and the vertical bars indicate the upper and lower 5 percentiles.

### Population dynamics and death in sub-MIC treatments

In the previous section we show that it is necessary to take death into account when computing mutation rate. For this, tracking population size (and thus net growth rate) during antibiotic treatment, as classically done by plating and counting colony forming units, is not sufficient. It is not possible to know whether a decreased net growth rate in the treatment compared to the untreated control is due to a purely bacteriostatic effect (i.e. the population grows more slowly, but without death) or to a bactericidal effect (i.e. the bacteria keep dividing, potentially at the same rate as without antibiotic, but also die). The first scenario will have no effect on the accumulation of mutants as a function of population size, while in the second scenario turnover implies a higher number of DNA replications and thus more mutants for a given population size, as explained above.

To disentangle these two effects, we designed a method allowing to compute growth rate and death rate simultaneously, using a segregative plasmid. The segregation dynamic permits to estimate the number of bacterial cell divisions. Combining this information with the change in population size permits to estimate growth rate and death rate, as explained in the materials and methods.

Our ultimate goal is to reliably estimate mutation rates of bacteria treated with sub-inhibitory doses of antimicrobials. To this end we quantify population dynamics and compute mutation rates toward a chosen neutral phenotype (resistance to rifampicin, conferred by substitutions in the gene rpoB) in populations exposed to sub-inhibitory doses of other antimicrobials. Our mutagenesis protocol is inspired by the standard fluctuation test with additional measurements of plasmid segregation to compute death rate, as detailed in the material and methods. The population dynamics are quantified as a combination of two variables: CFU at various time points (e.g. 0h, 3h, 6h and 24h after treatment starts), and relative death rate (compared to birth rate) between pairs of two successive time points. We represent these population dynamics in figure 3 for the chosen sub-inhibitory antimicrobial treatments. We use kanamycin at 3ug/mL, norfloxacin at 50ng/mL, and hydrogen peroxide (H_2_O_2_) at 1mM, allowing direct quantitative comparisons with the data from Kohanski and collaborators [28]. We find that for norfloxacin, there is a strong death rate in all phases of growth, and a strong impact of the treatment on final population size. For H_2_O_2_, death is only detectable in stationary phase and the treatment is mostly bacteriostatic during growth. For kanamycin, the dynamics are more complex, because an initially high death rate leads to a strong decline of population size during the first 6 hours of growth, followed by a recovery leading to a final population size close to the one reached in untreated controls. During this second phase of growth following the bottleneck at 6 hours, death rate is still substantial. This clearly shows that none of the three studied treatments are fully sub-lethal, and thus that the implicit hypothesis of no death made when using the standard methods of computation of mutation rate (as in Kohanski, DePristo, and Collins [28]) does not apply.

**Figure 3:**
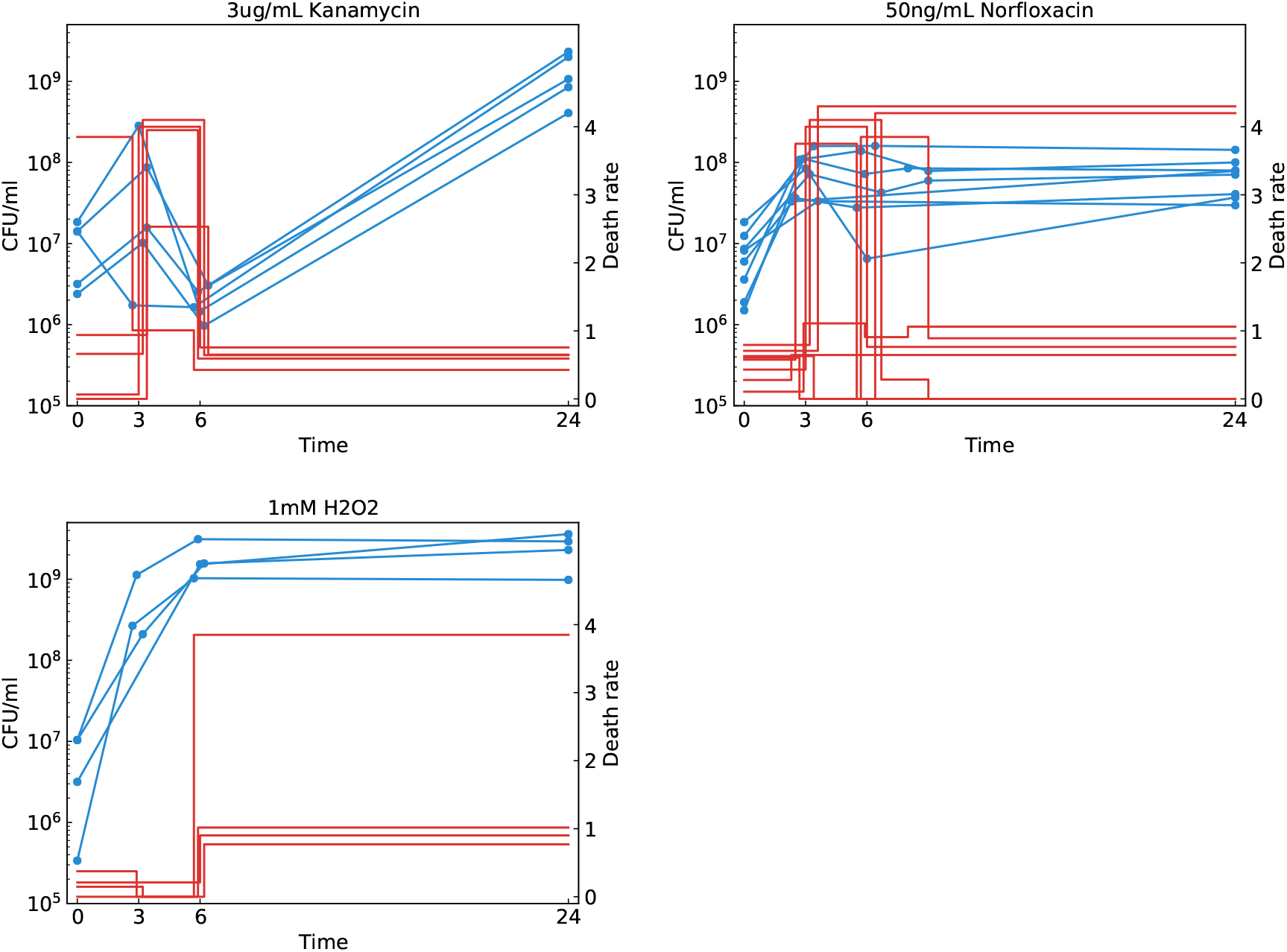
Growth and death dynamics of populations treated with sub-MIC antimicrobials. Each panel shows data for treatment with a different antimicrobial. Blue dots (left axis), joined by straight lines, represent population sizes measured by plating, expressed as colony forming units per mL of culture. Red lines (right axis) represent relative death rates, computed from plasmid segregation data, corresponding to the average number of death event per division event between two successive time points. Each line corresponds to a fully independent biological replicate, performed on a different day with a different batch of medium, and comprising at least 4 replicate populations. Time points 3 and 6 hours were horizontally shifted by a small offset value to avoid overlapping lines. Death rates higher than 4 were set to 4, plus or minus a small offset value to avoid overlapping lines.

### Mutation rate in sub-MIC treatments

We developed computational tools to quantify mutation rate taking into account the measured population dynamics, accounting for death. Our software, ATREYU (Approximate bayesian computing Tentative mutation Rate Estimator that You should Use) is described in the materials and methods. It takes as input any arbitrary population dynamics, described as a list of population sizes (i.e. CFU/mL for several time points) and an associated list of death rates between pairs of consecutive time points. This input is thus exactly what is shown in figure 3. We apply this method to analyse the results of our mutagenesis protocol, to quantify whether and by how much sub-inhibitory treatments with kanamycin, norfloxacin or hydrogen peroxide increase mutation rate. We show the effect of treatment on mutation rate in figure 4. We also plot the uncorrected mutation rate estimate assuming no death as would be obtained by methods such as FALCOR [26], bzRates [22], or rSalvador [59]. Clearly, not taking death into account leads to a strong overestimation of the mutation rate for both kanamycin and norfloxacin. In the case of kanamycin, correctly computing the mutation rate removes all signal for stress-induced mutagenesis. In the case of norfloxacin, this signal is strongly lowered, from a 14-fold to a 6-fold increase. For H_2_O_2_, the signal is less affected, which can be attributed to death rate being only significant in stationary phase. This confirms that neglecting death leads to a systematic over-estimation of mutation rates, and that taking into account the full population dynamics is necessary and leads to significantly different patterns depending on the antimicrobial and its effect on growth and death.

**Figure 4:**
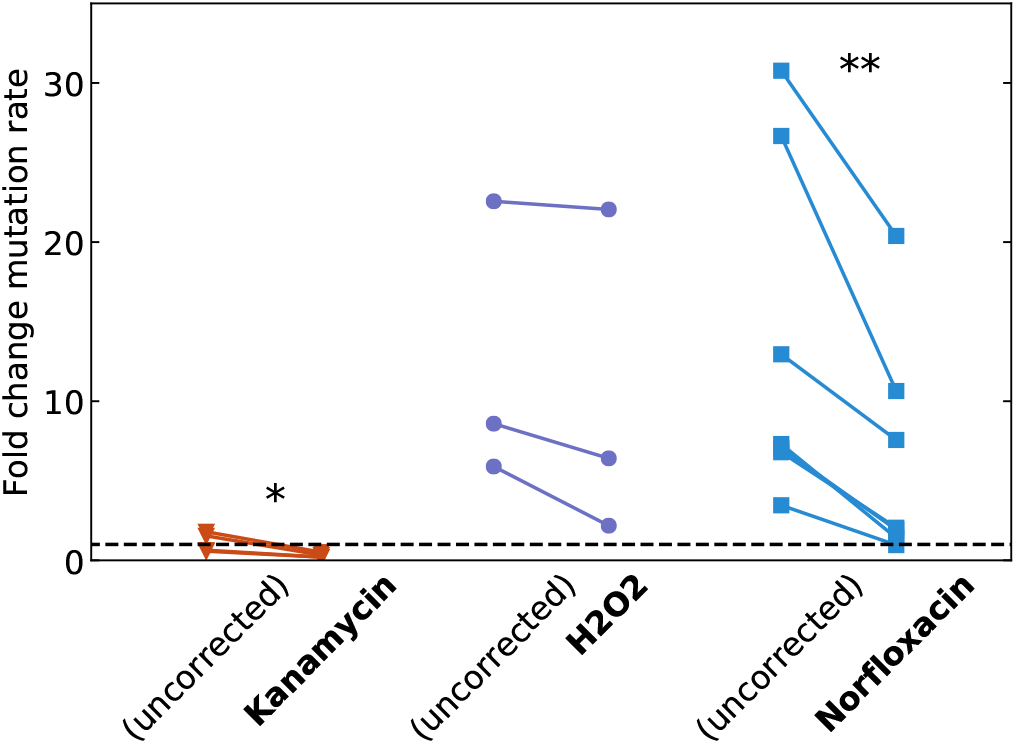
Change in mutation rate when treating with a sub-MIC dose of kanamycin, H_2_O_2_ or norfloxacin. The uncorrected mutation rate is the one that would be computed based on our data when death is not taken into account. Each point corresponds to a fully independent biological replicate comprising 24 parallel populations, from which population dynamics and mutation rates are estimated. All the computed mutation rates are normalised by the average mutation rate computed in absence of treatment. We performed a paired t-test to estimate whether the corrected mutation rate is significantly lower than the uncorrected one (* *p* < 0.05, ** *p* < 0.01).

### The link between evolvability and mutation rate depends on population dynamics

The quantification of mutation rate in different conditions is not sufficient to answer the question whether sub-inhibitory antibiotic treatments increase the likelihood of emergence of a resistant mutant and thus the probability of treatment failure. Indeed, mutation rate is expressed per DNA division, but, as we have shown in the previous section, antibiotic treatment may significantly change the number of susceptible cells and the number of replications that these cells have undergone. Intuitively, if a treatment multiplies mutation rate by 10 but divides population size by 100, it is not likely to lead to an increased genetic diversity. This intuition has also been given in Couce and Blázquez [11, figure 2, page 535], but has been largely ignored in the literature as it was not the main message of this review. Conversely, a treatment that does not affect mutation rate and only slightly affects carrying capacity but causes death and turnover may result in a significantly increased genetic diversity.

We first show the effect of sub-inhibitory treatment on final population size in figure 5. While H_2_O_2_ does not affect final population size, there is a strong effect of 1-2 orders of magnitude for norfloxacin, and a significant but smaller effect of around 50% reduction for kanamycin. This supports our intuition that at least for norfloxacin, the few-fold increase in mutation rate we report in the previous section is probably uncorrelated to any increase in genetic diversity.

**Figure 5:**
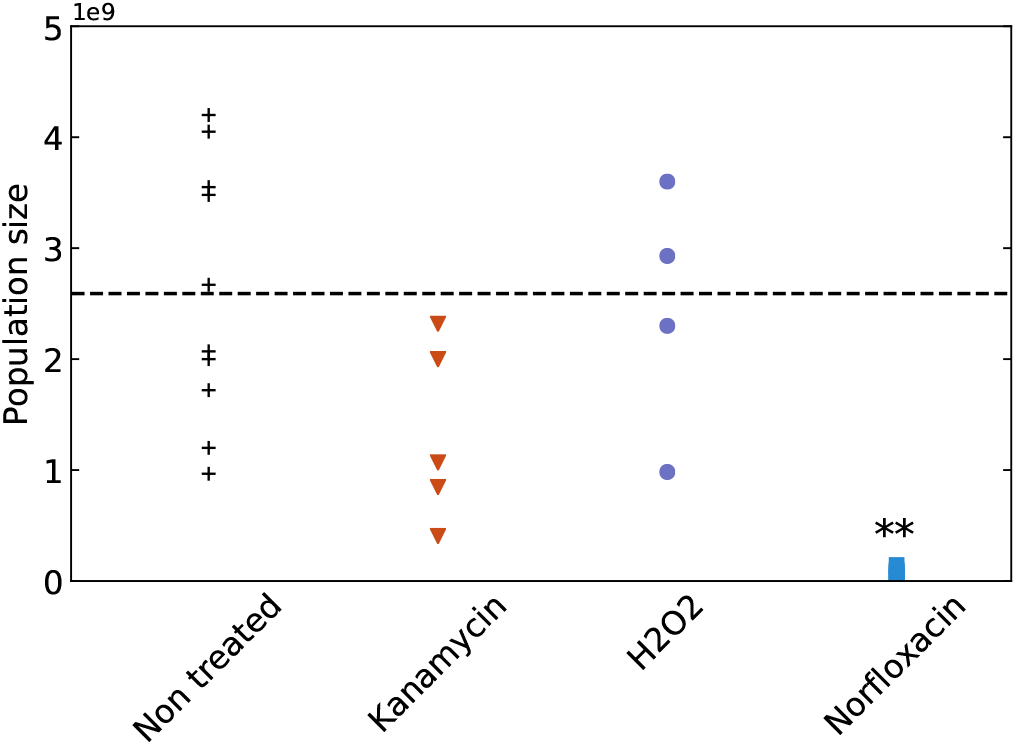
Population size reached after 24h in treated and untreated conditions. Each point corresponds to a fully independent biological replicate. For each of these biological replicates, average population size is estimated by plating appropriate dilutions of at least 6 replicate populations on non-selective medium. The dashed black line indicates the average size of the untreated populations. We performed an unpaired t-test to estimate whether treated population sizes are significantly different than untreated ones (** *p* < 0.01).

We expect the generation of genetic diversity to depend on (i) the number of cells alive, (ii) the population dynamics of these cells, and (iii) their mutation rate. Addressing the effect of stress on mutation rate as done in the previous section is necessary for a proper understanding of the bacterial stress response and of DNA repair mechanisms. Nevertheless mutation rate is not the relevant measure to understand the effect of stress on the generation of genetic diversity and thus on evolvability.

As a simple quantification of the generation of genetic diversity and thus of evolvability, we measure the number of mutants at a neutral locus, here the base-pair substitutions conferring resistance to rifampicin in the gene rpoB.

We plot in figure 6 the absolute number of rifampicin resistant mutants in the final population for all treatments and for untreated control. Evolvability is reduced by a few-fold by kanamycin treatment as expected, since this treatment decreases population size without increasing mutation rate. While norfloxacin and hydrogen peroxide both induce a small increase in mutation rate, they interestingly have strongly opposite effects on evolvability. Treatment with H2O2 increases evolvability by more than one order of magnitude, while treatment with norfloxacin reduces it by a similar amount. This is due to the very different effect these antimicrobials have on population dynamics: while H_2_O_2_ does not affect final population size, norfloxacin causes a strong decrease in population size due to both bactericidal and bacteriostatic effects.

**Figure 6:**
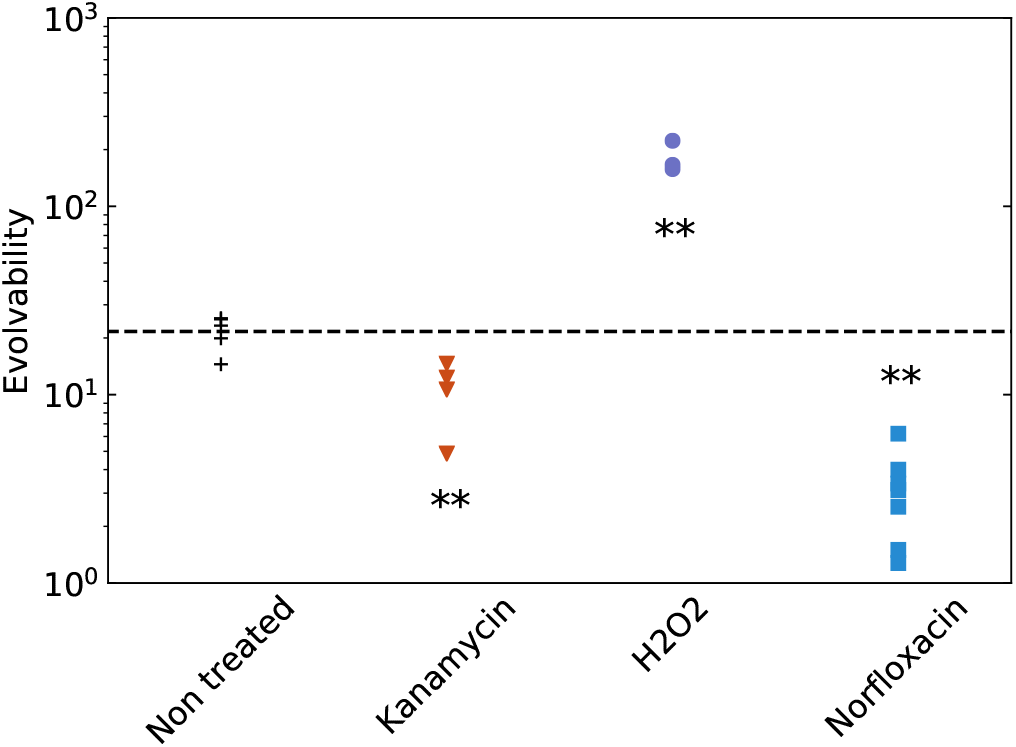
Evolvability of untreated and treated populations. For each treatment, we estimate evolvability as the number of rifampicin resistant mutants in the population after 24h of growth. The dashed black line represents the average evolvability of the untreated populations. Each point is a fully independent biological replicate comprising 24 replicate populations in which the number of rifampicin resistant mutants is averaged. We performed an unpaired t-test to estimate whether evolvability of treated populations is significantly different than this of untreated ones (** *p* < 0.01).

So independently of the question whether antibiotics increase mutation rate, we show that the sub-MIC treatments we studied do not in any way increase evolvability. Thus the standard rationale, that these sub-inhibitory treatments would increase the risk of emergence of resistance and treatment failure because of a higher generation of genetic diversity [5], does not hold.

This effect is largely due to a strong reduction of population size, which implies a loss in genetic diversity. Population size is however not the only factor affecting evolvability. We may also ask how much the measured turnover in our experiments contributes to evolvability. To answer this question, we simulate the same population dynamics as observed for each treatment, but without death: Each population reaches the same final population size as measured in our experiments, with the same mutation rate as computed, but with no death. This is similar to what would happen if the antibiotics only had a bacteriostatic effect. For each simulation, we quantify evolvability using the same measure as previously, i.e. the absolute number of mutants for our phenotype of interest in the final population. We compare this simulated evolvability without turnover with the actual measured evolvability in figure 7. For kanamycin and norfloxacin, turnover significantly increases evolvability by a few-fold.

**Figure 7:**
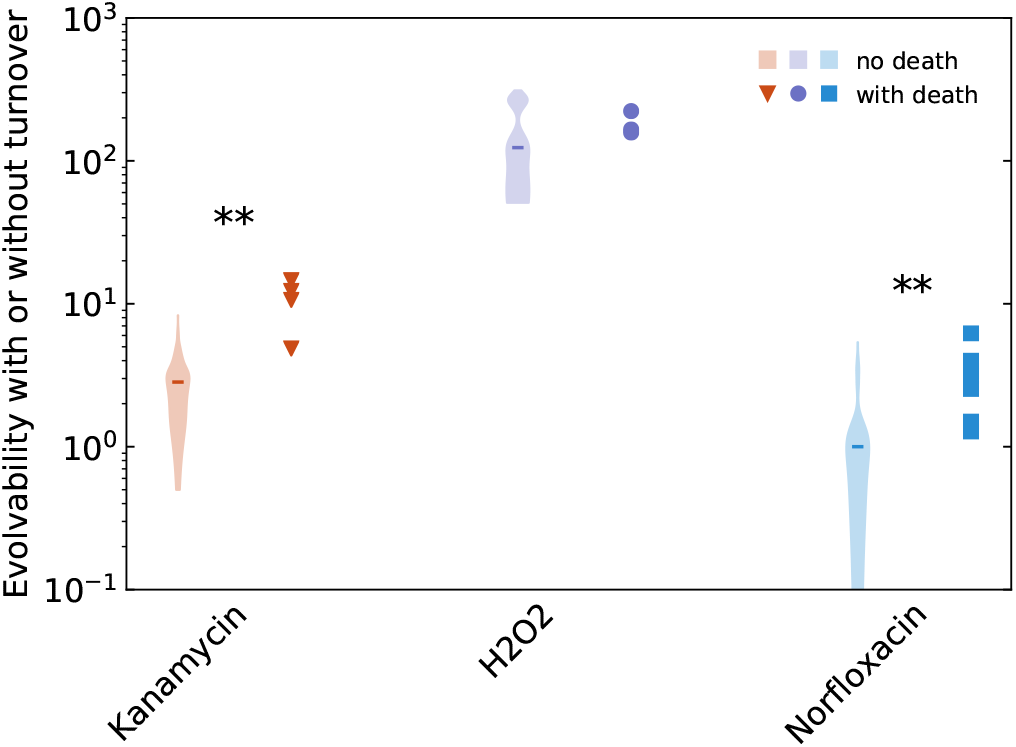
Contribution of turnover to evolvability. For each treatment, we plot the measured evolvability of the populations (red triangles, purple dots and blue squares) and the estimate that was obtained if the same population size was attained without cell death and with the same mutation rate (violin plots, 100 replicate simulations for each biological replicate). The median of all simulations for a given treatment is represented as an horizontal bar. For each treatment we performed an unpaired t-test to test whether the evolvability with death is different than the evolvability without death (** *p* < 0.01).

## Conclusions

In summary our results show that (1) mutation rate is systematically over-estimated in sub-inhibitory treatments because of death, (2) mutation rate is not a good predictor of the generation of genetic diversity or evolvability, (3) population size and turnover play a key role in evolvability, and (4) treatment with sub-inhibitory doses of norfloxacin or kanamycin significantly decreases evolvability, measured as the generation of genetic diversity at population scale. These results are in apparent disagreement with the conclusions of previous studies on antibiotic induced mutagenesis. This discrepancy is due to both miscalculation of mutation rates (because neglecting population dynamics) and misconceptions about the link between mutation rate and evolvability in these classical papers.

## Discussion

Understanding genetic and environmental control of evolvability is central for the understanding of microbial adaptation to constantly changing environments. Evolvability is defined as the capacity of a population to generate adaptive genetic diversity. This can be decomposed in two variables: the amount of genetic diversity generated by a population (often inaccurately attributed to the mutation or recombination rate only), and the fraction of this diversity that is adaptive. We are here interested in the former. Genetic control over the amount of generated genetic diversity has been studied for a long time in the field of mutation rate evolution [9, 51]. The existence of constitutive mutator alleles in bacteria has been discovered before the mechanisms of DNA replication [54], and the selection pressures leading to their transient increase in frequency have been elucidated through both theoretical and experimental studies [47, 37]. Observing the evolution and fixation of such mutator alleles from non-mutator lineages in a long-term evolution experiment [44] plausibly facilitated the acceptance of these theories. On the other hand, plastic, environment-dependent control over the generation of genetic diversity has been a controversial paradigm shift in bacterial evolution [20].

It has been proposed for a long time that various stresses can increase mutation rates in bacteria [46, 4], including those triggered by antimicrobial treatments [28, 3, 25, 41, 32]. Several molecular pathways have been shown to be implicated in this phenomenon, the emblematic one being the SOS response [38]. In this work we have shown that the effect of stress on mutation rate can not be computed properly with the existing tools, because the underlying mathematical models make the assumption that there is no stress, or more precisely, that the stress does not affect population dynamics. We develop experimental and computational tools to measure population dynamics and compute mutation rates under stress, and apply them to the question of mutagenesis due to antibiotic treatment. We have shown that the intuition that low doses of antibiotics are dangerous because they lead to a higher generation of diversity is based on a misinterpretation of valid experimental data for two reasons: (1) the increase in mutation rate is overestimated due to overly simplistic assumptions, and (2) a higher mutation rate does not lead to a higher genetic diversity if population dynamics are affected (e.g. if population size is reduced).

The question of emergence of resistance alleles due to low doses of antibiotics (reviewed by Andersson and Hughes [1]) can however not be entirely addressed by measuring the generation of genetic diversity. The study of evolution can be decomposed in two parts: generation of diversity, and natural selection acting on this diversity. While we have shown that treatment with a sub-inhibitory dose of norfloxacin does not increase but rather strongly decreases the amount of generated genetic diversity, it has also been reported that resistance alleles can be maintained and enriched by selection even at very low antibiotic concentration [24]. Such selection of pre-existent alleles may be a much more valid reason for concern about sub-inhibitory treatments. However, the literature is not as unanimous regarding bacteria residing within a patient with an immune system, rather than in a test tube [14]. It has for example been suggested that treating with a lower dose of antibiotics could slow down the selection of existing resistance alleles by decreasing their fitness advantage compared to the sensitive, wild type strain, without compromising the success of the treatment [40, 13]. Combining our results with these papers calls for a reevaluation of the evolution of antibiotic resistance at low doses of antibiotics.

The question of the potentially adverse effects of low doses of antibiotics has been of longstanding interest in the medical community, as is evidenced by the famous quote from Alexander Fleming’s Nobel lecture [18], “If you use penicillin, use enough”. However, given the time of this research (penicillin was discovered in 1928 and thus 15 before Luria & Delbrück), one should not be surprised that this often cited out-of-context advice relies on a rather Lamarckian reasoning in terms of educating rather than selecting for resistance:

> *Then there is the danger that the ignorant man may easily underdose himself and by exposing his microbes to non-lethal quantities of the drug make them resistant. Here is a hypothetical illustration. Mr. X. has a sore throat. He buys some penicillin and gives himself, not enough to kill the streptococci but enough to educate them to resist penicillin…* [18]

Our findings are also relevant outside of the context of evolution during antibiotic treatment.

As we mentioned, mutagenesis in bacteria under nutritional stress was a key development in the understanding of the bacterial stress response and DNA repair, with a recent regain of interest [29, 35]. Our experimental system can *a priori* not be applied to study starving bacteria, for two reasons: (1) our plasmid segregation method only gives sufficient signal in non-stationary populations, and (2) many of the observations on starvation-induced mutagenesis are dependent on the presence of some spatial structure (for example bacterial colonies on an agar plates [7, 46, 4]). In this second case the population dynamics become much more complex and are unlikely to be realistically approximated by a single relative death rate parameter. But the exact same questions remain to be elucidated in this field: how many cell divisions happen in these starving colonies? In batch cultures, is stationary phase really stationary, or is there some turnover and recycling as recently suggested [48]? And more importantly, where does death come from: is it an unavoidable, externally caused phenomenon; or is there an internal component, such as an altruistic programmed cell death [50], or just traits selected in other environments that give a maladaptation to certain stresses [23]?

Zooming out from evolutionary microbiology, mutagenesis research in bacteria shows an interesting parallel with recent advances in cancer research. For a given cell growth dynamics (organogenesis, from stem cells to an organised population of somatic cells), a higher mutation rate (expressed per cell division) will boost the accumulation of mutations and thus the risks of cancer. This increase in mutation rate can be genetic, such as in the case of hereditary nonpolyposis colon cancer caused by a deficiency of mismatch repair [6], or environmental, such as exposure to carcinogenic compounds [55, 27, 15]. All of this is now part of textbook science on cancer, and is similar to increase in mutation rate in bacterial populations due to genetic (mutator alleles [54]) or environmental factors (stress-induced [20] or stress-associated [34] mutagenesis).

Tomasetti and Vogelstein [52] recently reported that the number of stem cell divisions is a strong predictor of cancer risk per organ. This is in parallel with our findings, which show that the number of cell divisions is central to predict the generated genetic diversity in a population of cells. Tomasetti and Vogelstein caused a big controversy by concluding that cancers would thus mostly be due to “back luck” (i.e. unavoidable consequence of the large number of cell divisions) rather than to environmental factors (e.g. exposure to mutagenic chemicals). We show here that the generation of genetic diversity depends on both mutation rate and cell population dynamics, which is in line with many studies that have criticised the interpretation of the data made by Tomasetti and Vogelstein.

The challenge of understanding evolvability in bacterial population is thus strikingly similar to the one of understanding cancer, in the sense that the outcome depends on a complex interplay of extrinsic and intrinsic factors acting at different scales. In the case of bacteria, additional complexity stems from the fact that the same treatments may both impact the number of cell divisions (death and turnover) and the mutagenicity of each division. The picture is further complicated by the difficulty to disentangle the direct effects of the drug from the effects of the stress-response triggered by the drug. But fortunately, while separating and measuring each factor requires complex experimental methods and mathematical tools, measuring evolvability on neutral loci is simpler at least in bacteria. We hope that our study will encourage researchers in the field to question more not only the appropriateness of the tools they use for mutation rate estimation and the assumptions implicitely made by using these tools, but also the pertinence of the variable they choose to report.

## Materials and Methods

### Experimental setup

Our mutagenesis protocol is directly inspired by the one used by Kohanski and collaborators [28] (which is in turn similar to that of Luria and Delbrück [33]) with the inclusion of a segregative plasmid to compute death rate, as explained further below. A culture of *Escherichia coli MG1655* (with plasmid pAM34) is inoculated from a freezer stock and grown overnight in LB supplemented with 0.1mM IPTG and 100ug/mL of ampicillin (to ensure maintenance of pAM34). After the culture reaches stationary phase (at least 15 hours of growth), it is washed 3 times in normal saline (9g/L NaCl) to remove traces of IPTG, and then diluted 10,000 times in a 500mL baffled flask containing 50mL of LB (to maximize oxygenation). After 3.5 hours of growth, the culture is inoculated at ratio 1/3 in 24 culture tubes containing a total volume of 1mL of LB supplemented with one of the studied antimicrobials at sub-inhibitory concentration (3ug/mL kanamycin, 50ng/mL norfloxacin, 1mM hydrogen peroxide, or untreated control). After 24h of growth at 37°C, the cultures are plated at appropriate dilutions on 3 different LB agar medium: LB only to count the total number of bacteria (CFU), LB supplemented with 100ug/mL ampicillin + 0.1mM IPTG to count the number of bacteria bearing a copy of the segregative plasmid, and LB supplemented with 100ug/mL rifampicin to count the number of mutants toward the phenotype of interest. Additionally to this 24h time point, cultures are also plated on LB and LB + ampicillin + IPTG at intermediate time points (3h and 6h) to have a more accurate quantification of plasmid segregation dynamics and thus a better time resolution for the estimation of death rate. The plates are incubated between 15h and 24h for LB and LB ampicillin IPTG, and exactly 48h for LB rifampicin, before counting colonies.

### Measuring death using plasmid segregation

pAM34 is a colE1 derivative whose replication depends on a primer RNA put under the control of the inducible promoter pLac [21]. Under the presence of 0.1 – 1*mM* IPTG (non-metabolisable inducer of the lactose operon), the plasmid is stably maintained in every cell. When IPTG is removed from the growth medium, the plasmid is not replicating anymore, or not as fast as the cells divide, and thus is stochastically segregated at cell division. The decrease in plasmid frequency between two time points then permits to compute the number of bacterial cell divisions that occurred between these two time points. Combined with the change in population size, this allows to compute average death rate and growth rate between these two time points.

pAM34 also carries a betalactamase. The number of plasmid-bearing bacteria can thus be counted by plating an appropriate dilution of the culture on LB supplemented with 0.1mM IPTG (to ensure maintenance of the plasmid within colonies founded by a plasmid-bearing cell) and 100ug/mL ampicillin (to only permit growth of colonies founded by a plasmid-bearing cell). The total number of bacteria is determined by plating an appropriate dilution of the culture on LB.

Because mutational dynamics does not depend on time, we chose to compute relative death rate (ratio of death rate and growth rate as functions of time), which is the average number of death events per division event.

The link between plasmid segregation, death and number of divisions between two time points can be expressed mathematically as follows.

If we have:

- *F*: the frequency of cells bearing at least a copy of the plasmid, measured by plating on LB + ampicillin + IPTG
- *N*: the total number of cells, measured by plating on LB
- *res:* the rate of residual replication of the plasmid relative to the division rate in absence of IPTG
- *g*: the number of generations, i. e. the average number of duplications each genome present at final time did undergo
- *d*: the relative death rate (temporal death rate divided by temporal division rate)

The plasmid is diluted/segregated at each division following the equation

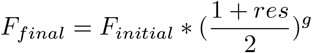

So we can estimate

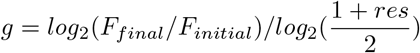

Without any death, we would have

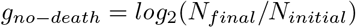

The difference between the true number of generations *g* computed from plasmid frequency and this number of generations *g_no–death_* computed based on the assumption that there is no death, permits to estimate relative death rate:

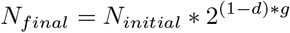

This yields

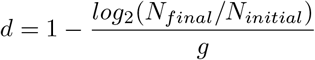

and thus

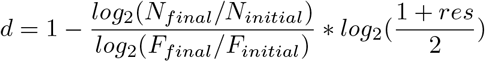

The only remaining free parameter to estimate is *res*, which is estimated by performing growth kinetics without antibiotic treatment (in LB medium) and thus without (or with negligible) death. We then have

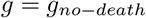

and thus

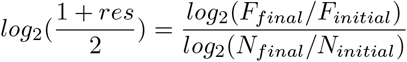

from which we can fit the value of the segregation parameter 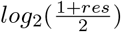 based on the values of *F* and *N* estimated by plating.

### Computing mutation rate taking death into account

Most modern measures of mutation rate rely on the same standard protocol, the fluctuation test [19], directly inspired by the Luria and Delbrück experiment [33]: several cultures are inoculated with a small population of non-mutant bacteria, are grown overnight and then plated on selective media (to count the number of mutants in the final population) and on non-selective media (to count the total number of bacteria in the final population). The number of mutants *r* in the final population (or rather its distribution over several replicate populations) is used to estimate the number *m* of mutational events happening during growth. One should note that these two numbers are not equivalent, because one mutational event can lead to several mutants in the final population if it happens early during growth, making this part of the computation complicated for intuition although good mathematical tools are available. The total number of bacteria *N* is assumed to be very close to the number of cell divisions (and thus the number of genome replications) because the initial number of bacteria is much smaller. Mutation rate can thus be estimated as *μ* = *m/N*.

The many existing software packages used to compute *m* from the observed distribution of *r* use an analytical expression of the probability generating function (pgf) of the number of mutants in the final population [30]. The only free parameter is the number of mutational events (equivalent to the value of the mutation rate per division when scaled with population size). This parameter is estimated from plating data using the maximum likelihood principle. The most used implementation of this idea is FALCOR (Fluctuation AnaLysis CalculatOR) [26], available on a webpage http://www.keshavsingh.org/protocols/FALCOR.html.

Other software packages implementing the same ideas have been developed more recently, including for example (r)Salvador [59] and bzRates [22], which also implement a few alternative assumptions such as fitness impact (cost or benefit) of the focal mutation, or a more accurate correction for plating efficiency than the one suggested by FALCOR [58].

However to this day, no available software permits to compute mutation rate when there is death. Some papers derived analytical expression of the pgf of the number of mutants in the final population in conditions where there is death [57], but this has to our knowledge never been applied to real data nor implemented in a software package. In theory such computations could easily be implemented in a tool similar to FALCOR (web server) or rSalvador (software package). However the basic assumption of the derived formula is that death rate is constant. This assumption is the price to pay for an analytical expression for the pgf, and is unfortunately not appropriate in our case given the observed death kinetics (see figure 3). On the other hand, given the computational power available today, we believe that analytical computations are not always necessary. In our case, while the measured population dynamics do not allow to derive an analytical expression of the pgf, it is straightforward to simulate many times such population dynamics with an arbitrary mutation rate, and to obtain an empirical distribution of the number of mutants. Running these simulations for any possible value of the mutation rate parameter then permits Bayesian inference: we look for the simulated mutation rate that gives the closest distribution to the one experimentally observed. Such methods are classically referred to as Approximate Bayesian Computing. We implemented such simulations in a Python program, and use this software as the heart of our data analysis.

## Acknowledgments

We acknowledge financial support from the Swiss National Science Foundation (SNF: 155866) and the European Research Council (ERC: PBDR 268540). We are grateful to Sandra Wotzka and Emma Slack for providing the plasmid pAM34.

